# A Phosphorylation Switch Governs KIF11’s Mechanical Output During Mitosis

**DOI:** 10.64898/2026.01.23.700875

**Authors:** Amila Šemić, Babu J N Reddy, Joseph M Muretta, Sarah E Vandal, Alex F Thompson, Cindy L Fonseca, Steven S Rosenfeld, Steven P Gross, Jason Stumpff

## Abstract

The kinesin-5 motor protein KIF11 is crucial for mitotic spindle assembly, driving the separation of spindle poles through microtubule sliding. Src-family kinases phosphorylate KIF11 at multiple tyrosine residues within its motor domain, but the mechanistic consequences of these modifications remain unclear. Here, we dissect the role of phosphorylation at Y211 using phospho-mimetic (Y211E) and non-phosphorylatable mutants (Y211F) in biochemical, biophysical, and cellular assays. Optical trapping and Förster resonance energy transfer (FRET) analyses reveal that Y211 phosphorylation slows neck-linker docking, reducing motor velocity and force generation under load. In human cells, Y211E expression impairs bipolar spindle formation and decreases spindle pole separation velocity, while Y211F shortens steady-state spindle length. Fluorescence recovery after photobleaching (FRAP) analyses show that Y211E accelerates motor turnover on spindle microtubules, consistent with the mutant motor’s heightened load sensitivity. Together, these findings support a model in which Src-mediated phosphorylation at Y211 acts as a rheostat to tune KIF11 mechanochemistry and spindle assembly dynamics, linking cancer-relevant kinase signaling to mitotic force generation.

## Introduction

Accurate chromosome segregation during mitosis depends on the formation of a bipolar spindle, a dynamic structure composed of microtubules (MTs) and numerous associated proteins. Spindle assembly requires the coordinated nucleation, organization, and sliding of MTs to establish a bipolar configuration that captures, aligns, and segregates paired sister chromatids accurately. This process depends on a balance of forces generated by molecular motor proteins and MT dynamics to maintain spindle length and tension while preventing collapse into a monopolar configuration. Among these motors, kinesin-5 family members play a central role by crosslinking and sliding antiparallel MTs apart to drive pole separation and stabilize spindle bipolarity. While kinesin-5 motors have been the subject of intense investigation, how their force production is regulated during mitosis to optimize spindle function remains incompletely understood.

Post-translational modifications of kinesin-5 motors have emerged as critical regulatory steps for optimizing their activity during mitosis. For example, phosphorylation of the C-terminal tail by cyclin-dependent kinase 1 (CDK1) or Aurora kinases has been shown to influence localization and function of *Xenopus laevis*, *Homo sapiens*, *Drosophila melanogaster*, *Saccharomyces cerevisiae*, and *C. elegans* kinesin-5 motors (1–7). CDK1 phosphorylation of the motor domains within the *S. cerevisiae* kinesin-5s Cin8 and Kip1 also regulates microtubule interaction and mitotic function (6, 8–10). Additionally, acetylation of human KIF11/Eg5 alters mechanochemical coupling to increase the motor’s resistance to load (11). These regulatory mechanisms highlight the importance of dynamic modification of kinesin-5 motors in adapting microtubule association and force production to the changing mechanical demands of spindle assembly.

Tyrosine phosphorylation of both the human and *Drosophila* kinesin-5 motors is also important for regulation during mitosis (12, 13). Src-family kinase phosphorylation of human KIF11 at tyrosine residues Y125, Y211, and Y231 has been observed both in vitro and in cells (13). Phospho-mimetic mutations, particularly Y211E, have been shown to reduce MT-stimulated ATPase activity and MT sliding velocity (13). Additionally, experiments utilizing LLC-PK1 cells suggest that optimal spindle assembly requires some level of KIF11 phosphorylation at Y211, as monopolar spindles form in Y211E-expressing cells, while disorganized spindles form in Y211F-expressing cells (13). However, it remains unclear how Src phosphorylation might affect KIF11’s motor mechanics and MT-binding properties in ways that could optimize spindle formation. This is an important question considering the contribution of abnormal Src-family kinases to oncogenic signaling (14).

In this study, we explore the biophysical consequences of Y211 phosphorylation on KIF11. Using a combination of optical trapping, Forster resonance energy transfer (FRET), live-cell imaging, and in vitro motility assays, we examine how phospho-mimetic and non-phosphorylatable mutations alter KIF11’s force generation, MT-binding dynamics, and spindle assembly function. By elucidating the mechanistic impact of KIF11 phosphorylation, our work provides insight into how regulation of KIF11 by Src-family kinases fine-tunes spindle dynamics to preserve genomic integrity.

## Results

### Phospho-mimetic KIF11 mutants display reduced velocities and force generation

To determine the effects of Src phosphorylation on KIF11 motor mechanics, KIF11 constructs that mimic phosphorylated motor (Y125E, Y211E, or Y231E) or unphosphorylated motor (wild type (WT) or Y211F) fused to the kinesin-1 coiled-coil and a C-terminal 6x-His-tag were expressed and purified from bacteria (Figure S1A). The use of this pseudo-dimer construct as a framework for testing the impact of Eg5/KIF11 regulation has been previously described (15). Purified recombinant proteins were attached to polystyrene beads via the His-tag, and KIF11 single molecule mechanics on purified MTs were measured using an optical trapping system that placed the motors under force while associated with a MT (Figure S1B). Phospho-mimetic KIF11 mutants (Y125E, Y211E, Y231E) exhibited reduced velocities and heightened force sensitivity compared to WT (Figure 1A-B). Among these, the Y211E mutant displayed the most significant decrease in velocity and force production with a peak force of 0.9 ± 0.03 pN compared to 2.2 ± 0.06 pN for WT. In contrast, the Y211F mutant closely resembled WT KIF11 in both velocity and peak force values.

**Figure 1.**
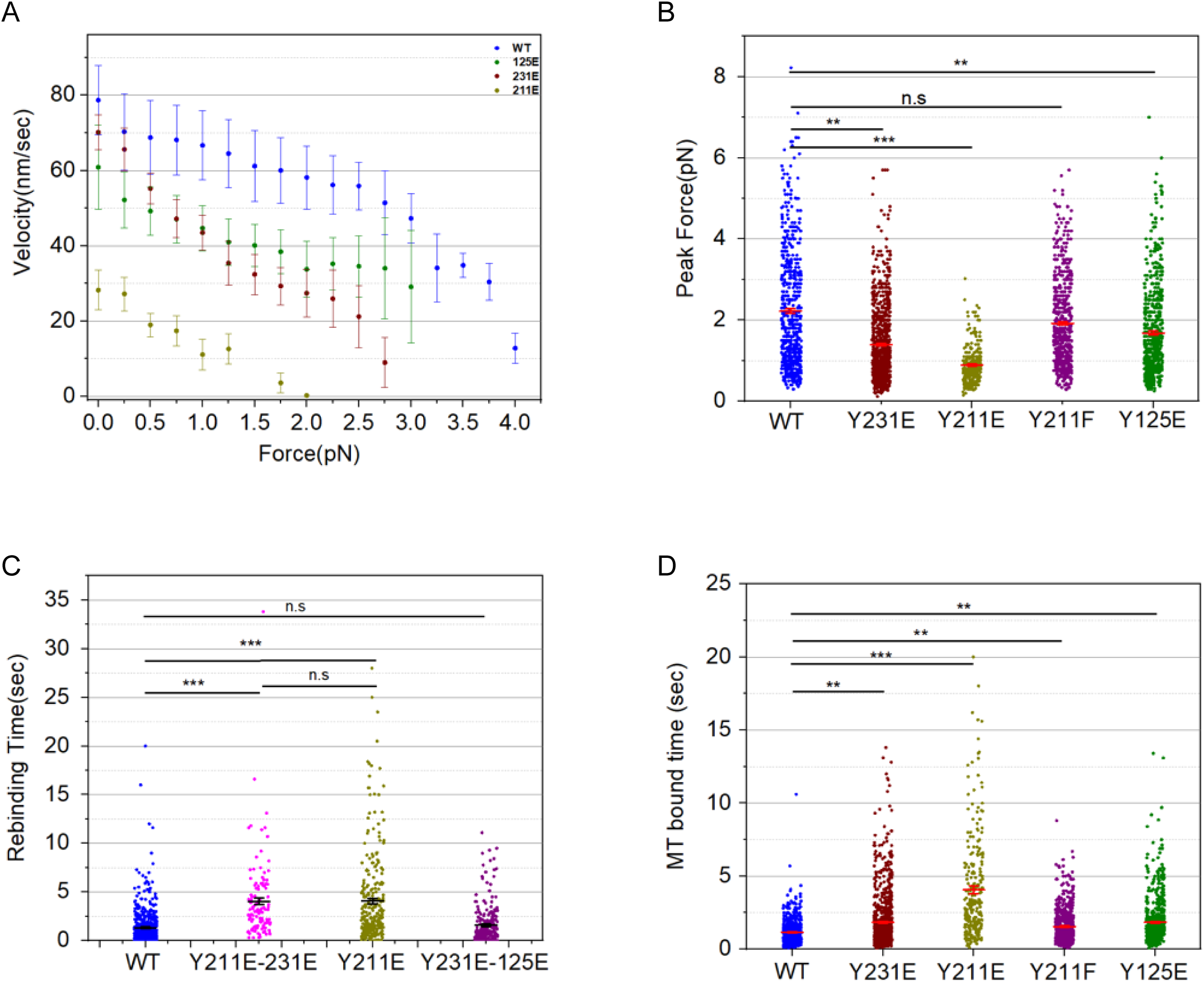
Phospho-mimetic mutations in KIF11 alter motor force production, velocity, and MT interaction dynamics. Force, velocity, and MT binding kinetics of single-molecule KIF11 pseudo-dimer constructs were measured using an optical trap. (A) Plot of force-velocity curves for the indicated constructs. N = 47, 65, 88 and 70 max force producing traces selected from 14, 9, 12 and 11 single-molecule beads of WT, 211E, 213E and 125E KIF11 motors. Error bars represent SEM for all traces at the given force bin. (B) Plot of peak forces measured for each of the indicated constructs. (C) Plot of MT bound times measured for each of the indicated constructs. Data points represent individual single-molecule measurements. (D) Plot of bead rebinding intervals measured for each of the indicated constructs. (B-C) Data points represent all single-molecule events of motor activity detected in the trap for the beads in 1 A. Mean ± SEM is indicated for each data set. Data were compared using student t-test (* p<0.05 *** p<0.001).

We also analyzed the effects of combining two phospho-mimetic mutations on KIF11 force production and found that the combination of Y211E/Y231E displayed very similar peak forces to the Y211E mutant alone (1.07 ± 0.04 pN compared to 0.9 ± 0.03 pN), further supporting the conclusion that Y211E has the greatest effect on force production (Figure S1C). On the other hand, the Y231E/Y125E double mutant displayed a peak force of 2.1 ± 0.07 pN, which is not significantly different from the 2.2 ± 0.06 pN force displayed by WT KIF11 (Figure S1A). The restored force production in the Y231E/Y125E double mutant compared to either single mutant suggests that each mutation may have long-distance conformational effects that are not necessarily additive. In any case, the data implicate Y211 as the most important site for altering force production.

To understand the binding kinetics of these constructs, both the duration of binding to a MT and the time required to rebind a MT after release were measured (Figure 1C-D). Binding durations for all mutants tested were significantly increased relative to WT KIF11, with the largest increase being displayed by Y211E (4.06 ± 0.27 sec compared to 1.2 ± 0.04 sec for WT) (Figure 1C). The rebinding interval was also increased relative to WT KIF11 for all constructs except Y231E, and the Y211E mutant again showed the largest change relative to WT (4.1 ± 0.27 sec compared to 1.33 ± 0.08 sec for WT) (Figure 1D). A double mutant combining the Y211E/Y231E mutations displayed binding durations and rebinding intervals similar to the Y211E mutant alone (Figure S1D-E). These differences indicate changes in MT-binding kinetics, where the phospho-mimetic mutations increase both the on-rate and off-rate of the motor, which could contribute to the spindle phenotypes previously observed in cells expressing KIF11 phospho-mutants. For example, the force generated by a group of Y211E motors could be lower than WT both because the single molecule force is lower for Y211E and because fewer molecules may be engaged with the MT. Taken together, these data suggest that phosphorylation of the KIF11 motor domain at Y125, Y211, and Y231 reduces motor force production, slows ATP hydrolysis, and alters the affinity of the motor for microtubules, with the charge change at Y211 having the greatest effect on these properties.

### Neck-linker docking is slowed by Y211E phospho-mimetic substitution

The observed effects of phospho-mimetic mutations on KIF11 force and velocity suggests that Src phosphorylation might alter the motor’s structural kinetics. Since the Y211E mutation caused the largest shift in force and velocity, we conducted FRET-based structural kinetics assays focusing on the Y211E mutant. Specifically, we used a FRET reporter assay that monitors neck-linker (NL) docking to evaluate the impact of Y211E on the mechanical step that couples ATP and microtubule binding with motor displacement and motility. A cysteine-lite human KIF11 motor domain construct with a pair of single reactive cysteine residues introduced at residues 30 and 228 were used to detect movement of the KIF11 neck-linker (KIF11NL) (Figure S1A). This monomeric construct was engineered with (KIF11NL-Y211E) or without (KIF11NL-WT) the phosphorylation-mimetic amino acid substitution Y211E and then labeled with a FRET donor [N-acetylaminoethyl-8-naphthylamine-1-sulfonate (AEDANS)] and non-fluorescent quencher [N-(4-dimethylamino-3,5-dinitrophenyl)maleimide (DDPM)]. Donor only (AEDANS)- or donor + quencher (AEDANS/DDPM)-labeled KIF11NL-WT or KIF11NL-Y211E proteins were bound with 4 molar stochiometric excess of MTs and incubated for 30 minutes at room temperature in the presence of apyrase to remove residual ADP and then mixed with 5 mM MgATP by stopped-flow at 25°C to initiate the ATP-driven neck-linker docking step as performed in previous studies (16).

Following stopped-flow rapid-mixing, we measured the total fluorescence emitted from the AEDANS donor, and the nanosecond timescale, time-resolved fluorescence decay every millisecond to plot the change in total fluorescence from MT complexes of KIF11NL-WT or KIF11NL-Y211E (Figure 2A). The time-resolved fluorescence decay waveforms were analyzed as described previously (16) modeling the change in waveform shape as a two-step transition between an undocked neck-linker with low-energy transfer between the probes, and a docked neck-linker with high-energy transfer between the probes, to determine how the mole-fraction of docked neck-linker changes with ATP-binding in each sample (Figure 2B). The mole fraction transients were fit by single exponential functions to quantify the impact of the Y211E amino-acid substitution on KIF11’s working stroke. Y211E markedly slows the neck-linker docking working-step of KIF11’s reaction cycle from an average of 22.8 s^-1^ to 7.3 s^-1^, consistent with single molecular mechanics (Figure 2C). This reduction in docking speed likely affects the coordination between ATP hydrolysis and motor displacement, decreasing force production and ATPase activity in the Y211E variant.

**Figure 2.**
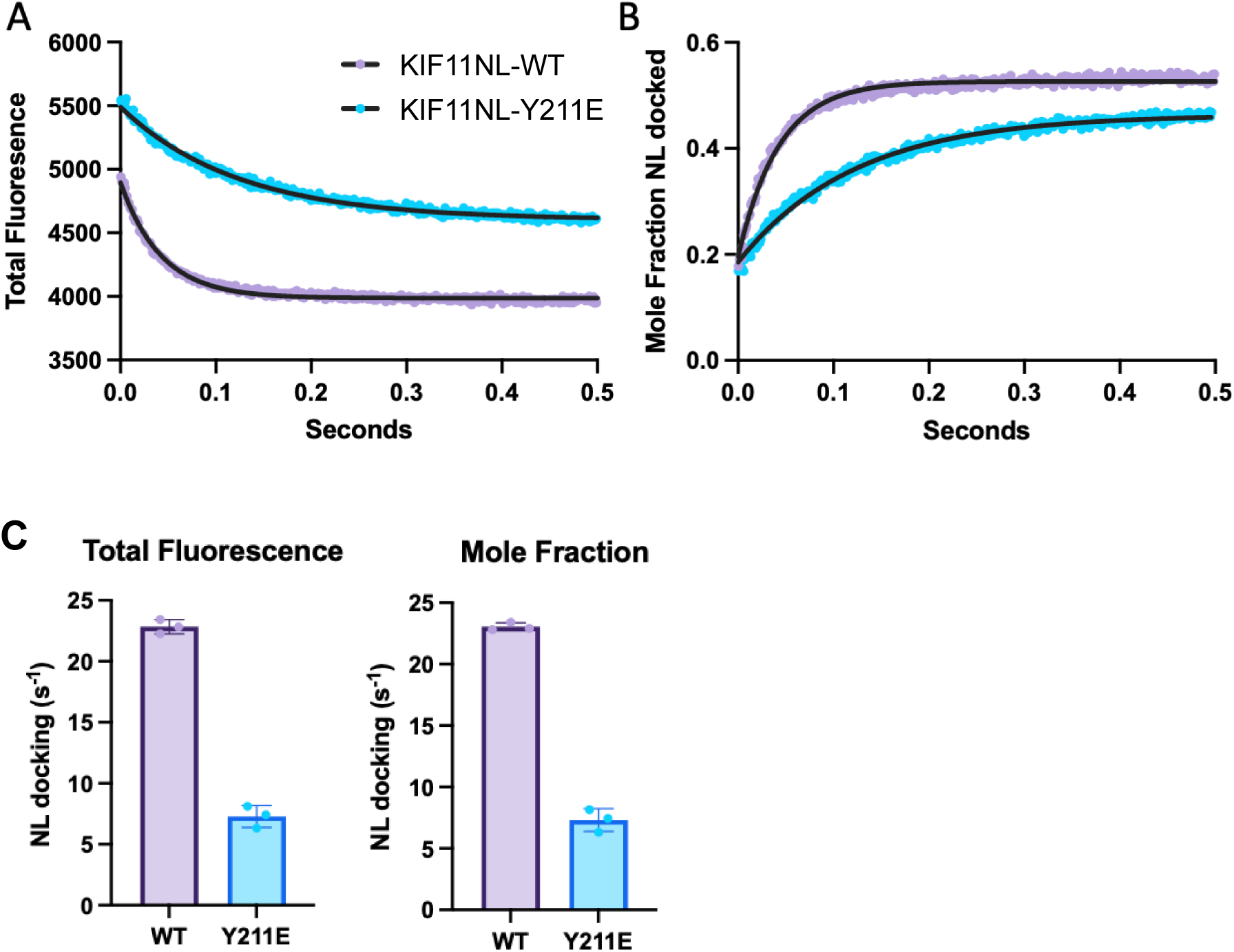
The Y211E phospho-mimetic mutation significantly slows KIF11’s neck-linker (NL) docking. NL-docking was measured by time-resolved FRET. (A) Total fluorescence transients of MT-bound AEDANS/DDPM-labeled KIF11NL-WT (magenta) or KIF11NL-Y211E (cyan) following mixing with 5 mM MgATP. (B) Mole-fraction transients obtained by analyzing time-resolved fluorescence decays to a two-state docked/undocked model. Single exponential fits to the transients (black lines). WT k = 22.43 s-1 (22.89 to 23.99 95% CI), Y211E k = 8.1 s-1 (7.917 to 8.298 95% CI). (C) Rate constants from N = 3 biological replicates measuring ATP-driven neck-linker docking measured by total fluorescence or model dependent time-resolved FRET mole fraction analysis.

### Y211 phospho-mimetic and phospho-null mutations alter mitotic spindle assembly

To determine how slowed neck-linker docking and reduced force production caused by phosphorylation of Y211 alters KIF11 function during mitosis in human cells, we generated stable, inducible RPE1 cell lines that expressed either siRNA-resistant KIF11 WT, Y211E, or Y211F with a C-terminal monomeric-Emerald-GFP tag (mEm). We confirmed that KIF11 siRNA treatment significantly depleted endogenous KIF11 (76% reduction in KIF11 immunofluorescent signal, Figure S2A) and that each stable cell line displayed a similar range of KIF11-mEm expression levels (Figure S2B). Mitosis was then analyzed in these stable RPE1 lines after treatment with either control or KIF11-targeting siRNAs and induction of the mEm-tagged transgenes. Analyses of fixed RPE1 cells confirmed that neither Y211E nor Y211F could fully rescue spindle assembly in KIF11-depleted cells. Expression of WT KIF11-mEm supported formation of bipolar spindles in 50% of cells (Figure 3A-B). We found that the fraction of WT KIF11-mEm cells that formed monopolar spindles expressed significantly lower levels of mEm than those that formed bipolar spindles (Figure S2B). In contrast, the majority of Y211E-mEm expressing cells formed monopolar spindles and none were observed to form bipolar spindles (100% monopolar), despite expressing mEm levels comparable to cells with bipolar spindles expressing WT KIF11-mEm. We also found that 35% of Y211F-mEm expressing cells formed bipolar spindles with 16% displaying short, disorganized spindles, consistent with prior studies in LLC-PK1 cells (Figure 3A-B) (13). Quantification of spindle length in the fraction of bipolar spindles in each experimental condition revealed that expression of Y211F led to significantly shorter spindles compared to controls regardless of whether endogenous KIF11 was depleted or not (Figure 3C). This suggests that inhibiting phosphorylation at Y211 has a dominant-negative effect on spindle length.

**Figure 3.**
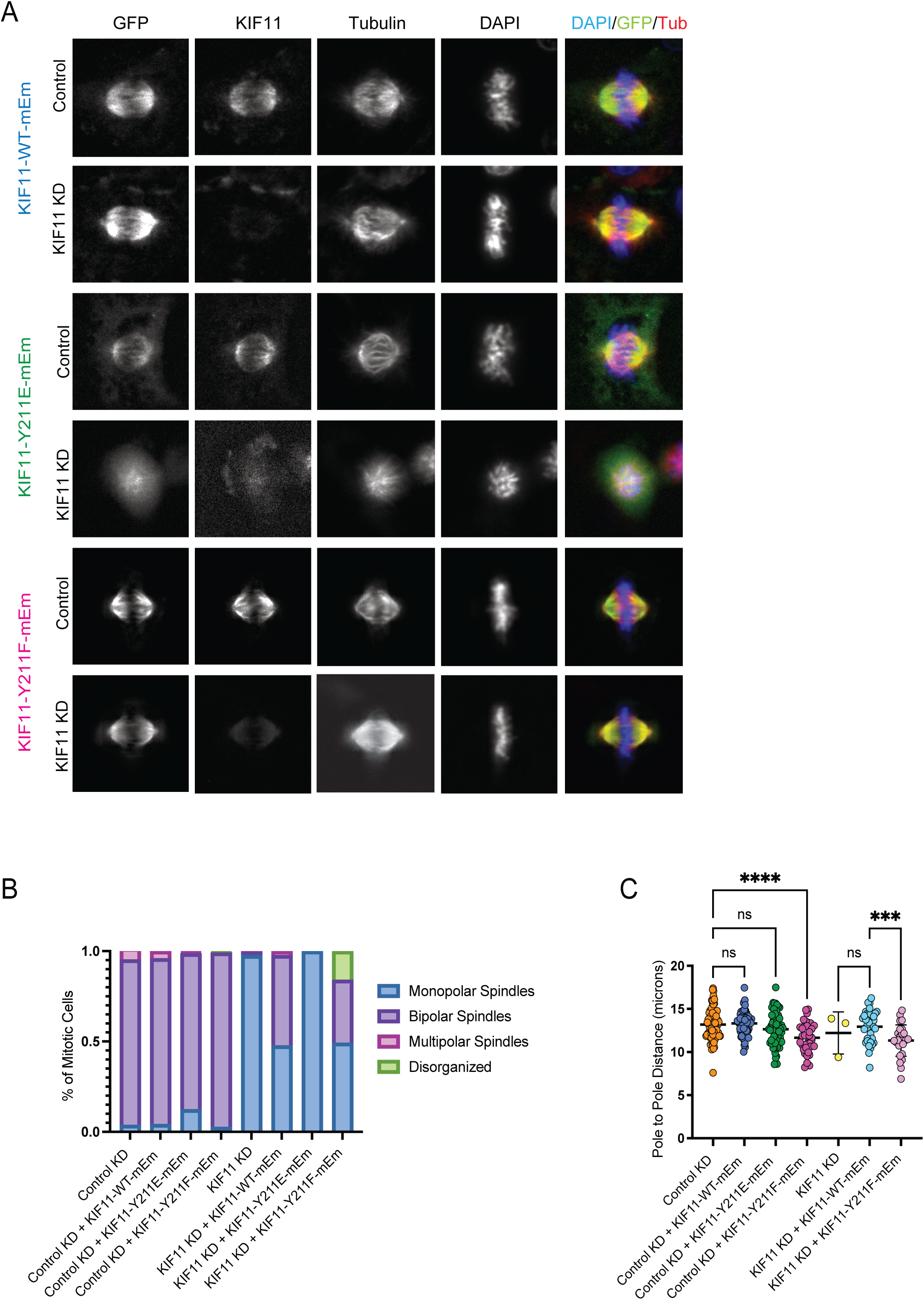
The KIF11 Y211E and Y211F mutations disrupt spindle formation in RPE1 cells. Spindle morphology was assessed in fixed RPE-1 cells expressing siRNA-resistant KIF11-mEm WT, the phospho-mimetic Y211E, or the non-phosphorylatable Y211F constructs. **(A)** Representative immunofluorescence images taken from each cell line, treated with either control or KIF11 siRNA. Cells were stained for GFP (green), DNA (DAPI blue), tubulin (red) or KIF11. Cells were fixed 24 hr after inducing expression of KIF11-mEm constructs with addition of doxycycline, and at least 24 hr incubation with siRNAs. **(B)** Mitotic cells were counted and categorized as having bipolar spindles (in metaphase), disorganized spindles, multipolar, or monopolar spindles. The percentage of cells with each spindle phenotype was determined for each condition. Cell counts for each cell line: **RPE1-KIF11-WT-mEM** - Control KD: 83; Control KD (+KIF11-WT-mEM): 73; KIF11 KD: 161; KIF11 KD (+KIF11-WT-mEM): 84; n = 3 experiments. **RPE1-KIF11-Y211E-mEm -** Control KD (+KIF11-Y211E-mEm): 70; KIF11 KD (+KIF11-Y211E-mEm): 247; n = 3 experiments. **KIF11-Y211F-mEm -** Control KD (+KIF11-Y211F-mEm): 38; KIF11 KD (+KIF11-Y211F-mEm): 63; n = 3 experiments. **(C)** The pole-to-pole distance was measured for bipolar spindles in each condition using the tubulin signal. Cell counts for each cell line: **RPE1-KIF11-WT-mEM** - Control KD: 75; Control KD (+KIF11-WT-mEM): 68; KIF11 KD: 3; KIF11 KD (+KIF11-WT-mEM): 41; n = 3 experiments. **RPE1-KIF11-Y211E-mEm –** Control KD (+KIF11-Y211E-mEm): 63; n = 3 experiments. **KIF11-Y211F-mEm -** Control KD (+KIF11-Y211F-mEm): 36; KIF11 KD (+KIF11-Y211F-mEm): 28; n = 3 experiments. Data were compared by running a one-way ANOVA with Šídák’s test for multiple comparisons: p-value style: <0.05 (*), <0.01 (**), <0.001 (***), and <0.0001 (****). If no significance is indicated, the result was not significant (>0.05). Scale bar: 5 µm.

To further investigate the effects of Y211 mutants on spindle assembly, we measured spindle assembly kinetics in cells expressing KIF11 WT-mEm, Y211E-mEm, or Y211F-mEm constructs. RPE1 cells were induced to express each construct and MTs were fluorescently labeled with SiR-tubulin. Cells were also treated with the KIF11 inhibitor monastrol to arrest mitotic cells with monopolar spindles in combination with control or KIF11 siRNAs. Following washout of monastrol, live cell imaging of pole separation dynamics in control siRNA-treated cells revealed that Y211E-mEm expressing cells exhibited a slower pole separation velocity than those expressing WT-mEm (Y211E: 0.39 ± 0.03 µm/min; WT: 0.74 ± 0.02 µm/min) yet ultimately reached similar maximum pole-to-pole distances (Y211E: 13.3 ± 0.2 µm; WT: 12.1 ± 0.4 µm) (Figure 4A-E; Movie S1). In contrast, KIF11-Y211F-mEm expressing cells exhibited similar pole separation velocities to those expressing KIF11 WT-mEm (0.61 ± 0.01 µm/min) but reached shorter maximum spindle lengths in both siRNA conditions (Control KD: 10.23 ± 0.3 µm; KIF11 KD: 9.0 ± 0.6 µm)(Figure 4A-E). These data suggest that the reduced force production and neck-linker docking of Y211E results in reduced pole separation velocity, even in the presence of endogenous KIF11. On the other hand, preventing phosphorylation of Y211 results in shorter spindles, potentially due to interference with endogenous KIF11 microtubule sliding.

**Figure 4.**
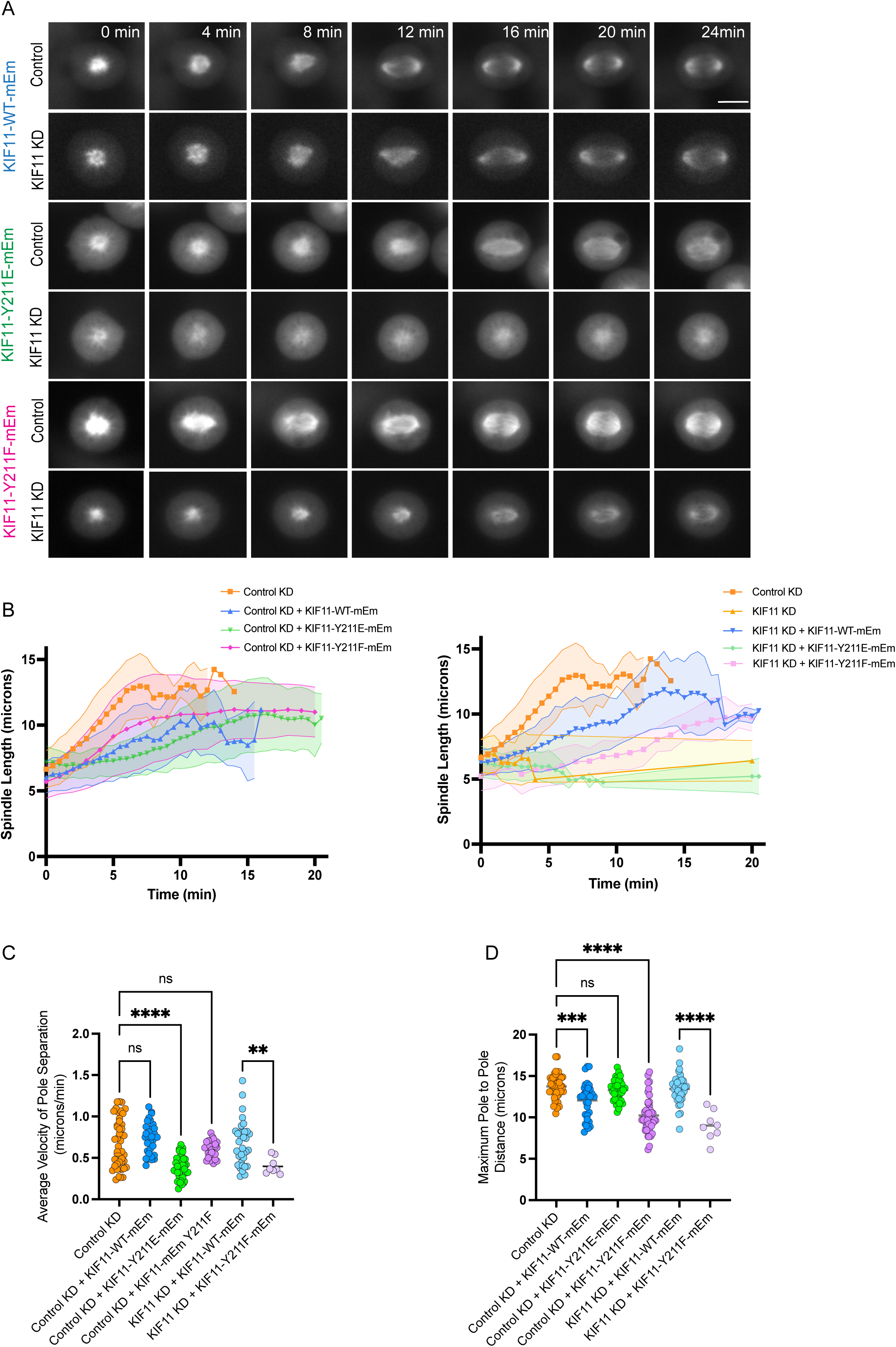
Expression of KIF11 Y211E slows spindle pole separation, while Y211F reduces maximum spindle length. Spindle polar separation was tracked in live movies of RPE1-KIF11-mEm expressing cells. Cells were induced to express KIF11-mEm constructs for 24 hr before live imaging with the addition of doxycycline and treated with control or KIF11 siRNA. Cells were also treated with monastrol overnight to inhibit KIF11 motors. This treatment collapses all spindles to monopoles and arrests cells in mitosis. Tubulin was labeled using SiR-tubulin several hours prior to imaging. Monastrol was washed out immediately before the start of imaging. Spindle pole separation was tracked using the SiR-tubulin signal after monastrol washout in individual cells at 30 sec intervals. **(A)** Representative stills of SiR-tubulin signal to show spindle pole separation after monastrol washout in each cell line, treated with either control or KIF11 siRNA. **(B)** The distance between spindle poles was tracked for cells in each line after monastrol washout, either in control siRNA (left) or KIF11 siRNA (right). **(C)** The average velocity of spindle pole separation was calculated for cells in each line after monastrol washout, either in control siRNA or KIF11 siRNA. Cells were excluded if spindle poles never began separating. **(D)** The maximum pole-to-pole distance achieved during spindle tracking. Cell counts for each cell line: **RPE1-KIF11-WT-mEM** - Control KD: 25; Control KD (+KIF11-WT-mEM): 27; KIF11 KD: 6; KIF11 KD (+KIF11-WT-mEM): 36; n = 3 experiments. **RPE1-KIF11-Y211E-mEm –** Control KD (+KIF11-Y211E-mEm): 31; n = 3 experiments. **KIF11-Y211F-mEm -** Control KD (+KIF11-Y211F-mEm): 31; KIF11 KD (+KIF11-Y211F-mEm): 8; n = 3 experiments. Data were compared by running a one-way ANOVA with Šídák’s test for multiple comparisons: p-value style: <0.05 (*), <0.01 (**), <0.001 (***), and <0.0001 (****). If no significance is indicated, the result was not significant (>0.05). Scale bar: 10 µm.

### KIF11-Y211E displays faster turnover from spindles during mitosis

To examine KIF11 kinetics within mitotic spindles, fluorescence recovery after photobleaching (FRAP) analyses were used to measure the turnover of KIF11 WT-mEM, Y211E-mEm, and Y211F-mEm constructs (Figure 5A-B; Movie S2). Stable RPE1 cell lines were induced to express each construct and treated with KIF11 siRNAs to increase the probability of measuring homotetramer behavior. KIF11-mEm fluorescence intensity was then measured before and after a 200 ms pulse with a 405 nm laser to bleach mEm in a circular region of interest with a 0.7 µm diameter. These analyses revealed that KIF11-Y211E-mEm exhibited faster turnover on spindle MTs (t_1/2_ =1.07 ± 0.06 sec) in KIF11 siRNA-treated cells when compared to KIF11 WT-mEm on either monopolar (t_1/2_ = 2.35 ± 0.37 sec) or bipolar spindles (t_1/2_ = 3.00 ± 0.36 sec) (Figure 5A-B). In contrast, Y211F-mEm displayed comparable turnover to WT-mEm in KIF11 siRNA treated cells (t_1/2_ = 2.26 ± 0.21 sec for bipolar spindles and t_1/2_ = 2.66 ± 0.31 sec for monopolar spindles). These data suggest that phosphorylation of Y211 increases KIF11 turnover on spindle MTs during mitosis. Interestingly, the turnover rate was similar between all KIF11-mEm constructs in FRAP experiments performed in interphase cells and overall, the rates were faster in interphase cells than on mitotic spindles (Figure S3). During interphase, KIF11 expression is reduced and is not thought to bind microtubules (2). Consistent with this, in KIF11-mEm constructs displayed cytoplasmic localization and did not appear to specifically localize to microtubules during interphase in RPE1 cells. These data suggest that the differences in turnover we observed for KIF11-mEm constructs on mitotic spindles are due to differences in how the constructs interact with microtubules, which is consistent with our biophysical data.

**Figure 5.**
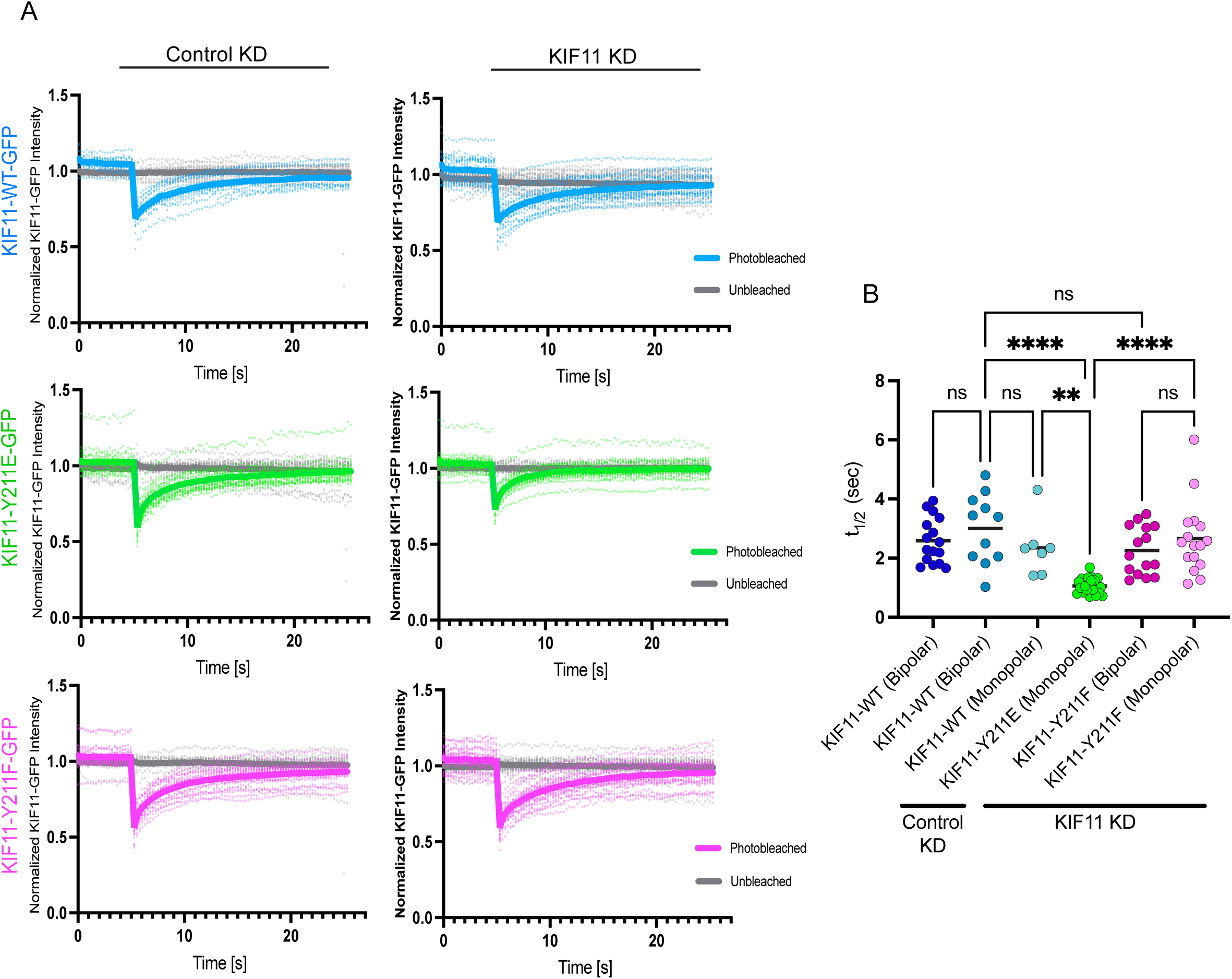
KIF11 Y211E displays faster turnover in mitotic spindles. KIF11-mEm construct turnover was measured in mitotic RPE1 cells using fluorescence recovery after photobleaching (FRAP). RPE1 cells were induced to express KIF11-mEm-WT, Y211E, or Y211F constructs with 24 hr doxycycline treatment, and either control or KIF11 siRNA. Using a 405 nm laser, cells were photobleached in a small region of interest on spindle microtubules, and fluorescence intensity recovery over the following 20 sec was measured. (A) Fluorescence intensity traces after photobleaching. Traces have been corrected for photobleaching due to image acquisition. (B) Half times (t1/2) calculated for each KIF11-mEm construct. Cell counts for each cell line: RPE1-KIF11-WT-mEM - Control KD: 16; KIF11 KD: 11 bipolar, 7 monopolar; n = 3 experiments. RPE1-KIF11-Y211E-mEm – KIF11 KD: 21, n = 3 experiments. KIF11-Y211F-mEm - KIF11 KD: 16 bipolar, 16 monopolar, n = 3 experiments. Data were compared by running a one-way ANOVA with Šídák’s test for multiple comparisons: p-value style: <0.05 (*), <0.01 (**), <0.001 (***), and <0.0001 (****). If no significance is indicated, the result was not significant (>0.05).

### Y211E does not impede MT sliding by WT KIF11

The slower pole separation velocities observed in cells expressing KIF11-Y211E in the presence of endogenous KIF11 suggests the phospho-mimetic motor may impede MT sliding of WT motor. To test this possibility directly, we performed MT gliding assays with mixtures of the same pseudo-dimer WT and Y211E constructs used for optical trapping studies (Figure S1A). Motors were attached to glass coverslips and MT movement across the surface was imaged and measured using TIRF microscopy. As expected, WT KIF11 exhibited significantly faster MT gliding velocity (13.4 ± 1.3 nm/s) compared to Y211E KIF11 (8.9 ± 2.7 nm/s) (Figure 6A). However, a significant reduction in MT gliding velocity was not observed when the two motor constructs were mixed, even at a 1:1 ratio, indicating that any MT sliding impedance by Y211E is negligible in mixed populations. This could be explained by Y211E rapidly detaching from the MT under load, consistent with our optical trapping and mitotic spindle FRAP results.

**Figure 6.**
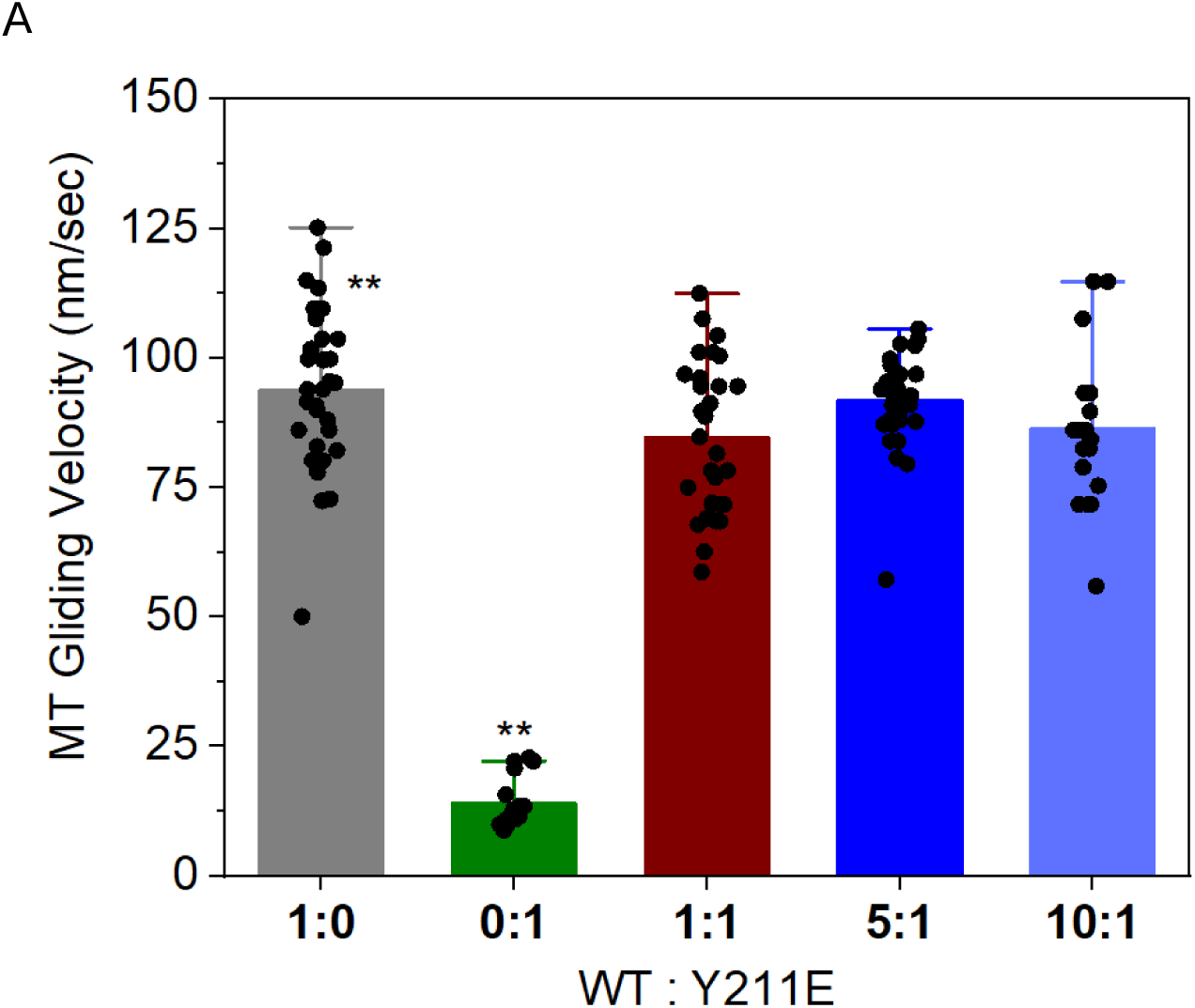
KIF11 Y211E does not impede microtubule sliding by WT KIF11. (A) Plot shows Cy-3 labeled MT gliding velocities in the presence of the indicated ratios of KIF11 WT and KIF11 Y211E pseudo-dimers. Data points represent the average velocities of individual motile MTs tracked for 200/1000 consecutive images. Number of MTs tracked, n = 32,17, 31, 33, 19 for WT:Y211E ratios of 1:0, 0:1, 1:1, 5:1, 10:1 respectively. Data were compared using a student t-test student (** p<0.01).

## Discussion

Our findings identify how Src phosphorylation of KIF11 tunes the mechanochemical performance of the motor and, in turn, regulates spindle assembly dynamics in human cells. Phospho-mimetic substitutions at three Src target sites within the KIF11 motor domain (Y125E, Y211E, Y231E) (13) reduce velocity and force output in single-molecule assays, with Y211E, the focus of this study, exerting the most pronounced effects. FRET-based structural kinetics analyses of Y211E indicate that this phospho-mimetic mutation slows neck-linker docking, which provides a mechanistic explanation for reduced stepping rate and force generation. In cells, these biochemical changes manifest as alterations in spindle assembly and motor turnover on spindle MTs. Notably, Y211E does not measurably impede MT gliding by WT KIF11 in gliding filament assays, consistent with a model in which the phospho-mimetic motor detaches rapidly under load rather than acting as a brake for MT sliding within the spindle. This interpretation is further supported by the increased turnover of Y211E from spindles in mitotic cells. Taken together, our data suggest that phosphorylation of Y211 functions as a rheostat that modulates KIF11’s working stroke and load tolerance to ensure proper spindle architecture and dynamics.

### Phosphorylation at Y211 slows the working step that couples ATP binding to forward displacement

Our optical trapping results demonstrate that phospho-mimetic KIF11 motors exhibit reduced velocities and heightened force sensitivity compared to WT, with Y211E demonstrating the largest changes in both metrics. Stopped-flow FRET experiments indicate that these changes in load-dependency are mechanistically linked to the slow neck-linker docking observed for Y211E. Neck-linker docking is the core conformational change that couples ATP binding and microtubule engagement to forward movement of the motor’s catalytic core (17). Thus, a slowed docking step is expected to decrease stepping frequency and reduce peak force by increasing the probability of motor detachment from the MT under load. This agreement between the FRET kinetics and single-molecule measurements supports a direct causal link between phosphorylation at Y211 and changes to the structural sequence that defines the kinesin working stroke.

Although Y125E and Y231E also diminish velocity and force, the larger effect of Y211E suggests this site is particularly important for the coupling between nucleotide state and neck-linker movement. The phospho-null Y211F, in contrast, largely resembles WT in optical trapping assays, indicating that the presence of a hydroxyl group to accept a phosphate at Y211 may be important for tuning motor performance rather than being essential for baseline mechanics. Taken together, our data are consistent with the idea that phosphorylation near the neck-linker interface can modulate the timing and stability of the docked state, thereby regulating load tolerance and processivity.

### Cellular consequences of disrupted Y211 phosphorylation

In KIF11-depleted RPE1 cells, neither Y211E nor Y211F fully rescues spindle assembly, whereas WT KIF11-mEmerald supports bipolar spindle formation in a substantial fraction of cells. Importantly, cells expressing Y211E form predominantly monopolar spindles despite comparable expression levels to WT-expressing cells that form bipolar spindles.

This phenotype is consistent with findings from LLC-PK1 cells (13), and aligns with the motor’s reduced velocity and force sensitivity under load, which would compromise crosslinking-driven sliding of antiparallel MTs required to separate poles. Y211F, while more WT-like in single-molecule assays, yields a distinct cellular phenotype: shorter, disorganized spindles even when endogenous KIF11 is present (Figure 3C). This suggests that preventing phosphorylation at Y211 dampens the ability of the spindle to expand to its normal steady-state length, pointing to a physiological role for dynamic phosphorylation in fine-tuning spindle architecture.

Live-cell imaging during monastrol washout corroborates these interpretations. In the presence of endogenous KIF11, Y211E expression reduces pole separation velocity but permits attainment of near-normal maximal pole-to-pole distances. We interpret this as a kinetic impairment, characterized by slower sliding under opposing load, rather than an absolute inability to reach longer lengths when the endogenous motor can compensate over time. In contrast, Y211F yields normal separation velocities but shorter maximal spindle lengths, consistent with a dominant effect on steady-state length control. This could potentially occur by motor overcrowding and impedance, leading to an alteration in the forces produced by KIF11 ensembles.

FRAP analyses further indicate that phosphorylation influences KIF11 MT engagement dynamics. KIF11-Y211E displays faster turnover than WT on spindle MTs, which we attribute to its heightened load sensitivity and reduced stability of the strongly bound state arising from slowed neck-linker docking. In contrast, KIF11-Y211F has comparable turnover to KIF11-WT on both bipolar and monopolar spindles. Taken together, these data suggest that phosphorylation at Y211 increases detachment rate or reduces dwell time on spindle MTs, thereby modulating motor availability and ensemble behavior during mitosis.

### Y211E does not impede MT sliding by WT KIF11

A key question raised by the slow pole-separation velocities measured in Y211E expressing cells is whether Y211E acts as a brake when co-expressed with WT KIF11. In MT gliding assays with mixtures of WT and Y211E dimeric motors, we did not observe a significant reduction in MT gliding velocity, even at 1:1 ratios. This implies that the phospho-mimetic motor does not function as a dominant roadblock when motors operate under loads in the geometry of gliding assays. This lack of a dominant-negative effect also aligns with our FRAP and optical trapping data, which indicate that Y211E detaches rapidly under load. Together, these results further support the idea that KIF11 motors phosphorylated at Y211 minimally contribute to ensemble force production and do not impose friction on WT motors.

### Model: Y211 phosphorylation as a rheostat for KIF11 ensemble kinetics

Integration of our biochemical, biophysical, and cellular data supports a model in which Src phosphorylation at Y211 adjusts the contribution of KIF11 motors to force generation within the spindle. Src phosphorylation at Y211 slows neck-linker docking, reducing the probability that the motor completes its working stroke before detachment under load. This reduced load tolerance and faster turnover, in turn, decreases the effective force per motor and the number of motors engaged in MT sliding at any given time. At the ensemble level, too much phosphorylation manifests as slower pole separation (Y211E), while preventing phosphorylation disrupts the capacity to reach normal, steady-state spindle lengths (Y211F). In this framework, we propose that phosphorylation of Y211 provides a tunable control point that could adjust KIF11’s contribution to spindle force generation, potentially in a spatial or temporal manner. Such a mechanism would also allow cells to modulate spindle kinetics and architecture without fully switching KIF11 “on” or “off.”

Allosteric inhibitors (e.g., monastrol) and domain-specific perturbations of KIF11/Eg5 have long demonstrated the sensitivity of spindle assembly to Eg5’s force generation, load tolerance, and processivity (18–21). Our results extend this paradigm by implicating Src-mediated phosphorylation at a specific tyrosine within the motor domain as a means to modulate the core mechanochemical step, neck-linker docking, that underlies forward displacement. Prior reports have described regulation of mitotic kinesins by phosphorylation and other post-translational modifications, often affecting MT affinity, autoinhibition, and cargo interactions (22). The Y211 site appears to influence the timing and stability of the docked neck-linker state, providing a direct link between kinase signaling and the mechanical output of a mitotic motor.

Beyond basic cell biology, these findings have potential relevance for contexts in which spindle dynamics are perturbed, such as in cancer cells with altered kinase signaling. Fine-tuning motor load tolerance and turnover via phosphorylation could contribute to adaptive changes in spindle assembly timing, robustness, and architecture. Conversely, manipulating Y211 phosphorylation, pharmacologically or genetically, could modulate sensitivity to Eg5 inhibitors and potentially influence mitotic progression in specific cellular states (13).

Future work should explore additional structural and kinetic insights of KIF11 phosphorylation by Src, such as defining when and where phosphorylation occurs within the spindle and across different phases of mitosis and how Src exhibits this spatial and temporal control in both normal and tumor cells. Additionally, structural analyses could define precisely how phosphorylation alters the neck-linker interface and nucleotide-dependent conformational transitions. These additional details would provide clarity regarding our working model that phosphorylation at Y211 fine-tunes KIF11 motor mechanics to promote optimal spindle formation and maintenance.

## Methods

### DNA constructs

All protein expression constructs were engineered into the pET vector system using Gibson assembly master mix (New England Biolabs #E2611L) with the vector linearized by restriction digestion. The WT KIF11 pseudo-dimer and WT motor domain NL FRET constructs are described in our previous publications (16). For plasmids containing the Y211E amino acid substitution, the KIF11 reading frames were synthesized as gBlocks (Integrated DNA Technologies) codon optimized for expression in *Escherichia coli* or amplified by PCR from existing plasmids. All constructs were validated by Sangar sequencing prior to transformation into the NiCo21(DE3) *Escherichia coli.* Strain (New England Biolabs #C2529H) for expression.

### Protein Expression and purification

The KIF11 pseudo-dimer was used for single molecule experiments. Sequence-verified expression plasmids were transformed into the NiCo21(DE3) *Escherichia coli.* and were grown to an OD600 of 1.6 then expressed by IPTG induction for 12 h at 18 °C. Cells were harvested by centrifugation, and the harvested pellets were resuspended in lysis buffer (50 mM HEPES, pH 7.5, 500 mM NaCl, 10 mM imidazole, 4 mM MgCl2, 5% Sucrose, 0.2 mM ATP, 5 mM 2-mercaptoethanol) and disrupted by sonication in the presence of protease inhibitors. The lysate was clarified by ultra-centrifugation at 185,000 RCF for 40 minutes and the clarified supernatant subjected to gravity-flow affinity purification using HisPur NiNTA resin (Thermo Fisher Scientific #88222). The resin was washed with 10-20 column volumes of wash buffer (50 mM HEPES, pH 7.5, 150 mM NaCl, 25 mM imidazole, 4 mM MgCl2, 5% Sucrose, 0.2 mM ATP, 5 mM BME). After washing, the protein was eluted in 5 column volumes of elution buffer (50 mM HEPES, pH 7.5, 150 mM NaCl, 250 mM imidazole, 4 mM MgCl2, 5% Sucrose, 0.2 mM ATP, 5 mM BME). The eluted protein was concentrated by centrifugation, clarified by centrifugation at 20,000 RCF for 30 minutes, and subjected to size exclusion chromatography on a Hi-Prep 16/6- Sephacryl-S200 HR column. Fractions containing the KIF11-psuedo-dimer were pooled and snap frozen in liquid Nitrogen (50 mM HEPES, pH 7.5, 150 mM NaCl, 4 mM MgCl2, 5% Sucrose, 0.1 mM ATP, 1 mM DTT) and stored at −80 °C.

### FRET Constructs

Monomeric motor domain constructs were engineered as described in our previous publications (16). Previously generated human KIF11NL constructs with reactive cysteines at positions 256 and 356 were engineered with the Y211E amino acid substitution and expressed and purified as previously described (16). Protein constructs were labeled with 1:8 molar stoichiometry of 1,5 IAEDANS (Thermo Fisher Scientific) to protein at 4°C overnight in ATPase buffer: 50 mM potassium acetate, 25 mM HEPES, 5 mM Mg Acetate, 1 mM EGTA, 0.5 mM TCEP, pH 7.50. Unreacted probe was removed by gel filtration on Sephadex G25 pre- packed columns (Pharmacia, PD10). Samples were then labeled with a 10-fold molar excess of DDPM over protein in ATPase buffer without the TCEP overnight at 4°C. The ATPase activity of these constructs was determined in ATPase buffer by measuring phosphate production in the presence of a minimum of a 5-fold molar excess of microtubules.

### Stopped-flow FRET experiments

Neck-linker docking in response to ATP binding to motor domain-microtubule complexes was measured using a time-resolved fluorescence equipped stopped-flow device described in our previous publications (16). Transient changes in the total AEDANS fluorescence and the nanosecond-scale AEDANS time-resolved fluorescent decay curves were analyzed as described in that work to determine the mole fraction of docked and undocked neck-linker in the motor domain constructs.

### Single-molecule in vitro motility assay

The optical trapping setup was assembled on an inverted Nikon TE200 microscope using a 980 nm single-mode fiber-coupled diode laser (EM4 Inc) and additional optical components. For all the single motor experiments with KIF11 (KIF11 head fused to Kinesin-1 coiled-coil tail and a His tag at the C-terminus) and its variants, the trapping laser power was set constant to achieve a trap stiffness (k_trap_) of approximately 0.045 pN nm^−1^ with a 0.56 μm polystyrene bead (anti-His antibody coated on streptavidin beads from Spherotech).

Sample chambers were constructed using polylysine-coated 0.17-mm thick coverslips and 50 μm double-sided adhesive tape as reported previously for in-vitro kinesin-1 motility experiments (23). Single motor experiments were carried out in the motility buffer (80 mM PIPES @ pH 6.9, 50 mM CH3COOK, 4 mM MgSO4, 1 mM DTT, 1 mM EGTA, 10 μM Taxol, 1 mg mL^−1^ casein) filtered with 100 nm centrifugal filter (Millipore Ultrafree-MC-VV) to minimize dust particles from interfering with motor binding in the laser trap. The motility buffer was supplemented with 2 mM ATP and an oxygen-scavenging system for measurements. Pseudo-dimer KIF11-WT or KIF11-mutant-coated polystyrene beads were prepared immediately before the measurements. Depending on the experiment, the purified KIF11 motors or their mutant versions were diluted to approximately 20 nM and mixed with approximately 1 pM of biotinylated penta-His-antibody-conjugated streptavidin beads (stock stored at 4°C), yielding a binding fraction of 10–20%. The incubation of beads and motors, approximately 50 μL, was carried out at room temperature for 10 minutes. Before injecting the mixture, the sample chamber with pre-assembled microtubules was washed with 50 μL of motility buffer. The video and high-resolution position data of bead binding events were recorded using a DIC microscope with a photodiode positioned in the back-focal plane of the condenser. This was accomplished with NI-LabVIEW software and digitization cards.

### Scoring Criteria for Binding Events

Data collection for single-molecule kinesin motility experiments was performed as documented in prior studies (11). Force data were recorded at 3 kHz using a position-sensitive detector and filtered through a 40-point FFT function. A binding event was confirmed when the position signal exceeded a threshold of 15 nm, lasting a minimum of 10 msec to differentiate true binding from noise. The motor detachment was recognized by sudden slope changes in the force signal. "Re-binding time" was defined as the duration between detachment and re-binding events detected by the threshold. In contrast, bound time was the motor activity period during which the force/position signal remained above the threshold before detachment. Force velocity profiles were estimated from the selected stall events for each bead as described previously (11).

### Cell culture

RPE1 acceptor cells for High-efficiency low-background (HILO) recombination mediated cassette exchange (RMCE) were described previously (24). Cells were cultured at 37°C with 5% CO_2_ in MEM-α medium (Life Technologies, 12561072) supplemented with 10% Fetal Bovine Serum (FBS; Life Technologies, 16000044). RPE1 acceptor cells for recombination were maintained in 10% blasticidin (ThermoFisher Scientific, R21001). siRNA transfections were performed using RNAiMax (Life Technologies) according to manufacturer’s recommendations. For siRNA transfections in a 24-well format, approximately 20,000 cells in 500 µL MEM-alpha medium with 10% FBS were treated with 15 pmol siRNA and 1.5 µL RNAiMax (Life Technologies) precomplexed for 10-15 minutes in 50 µL OptiMeM (Life Technologies). Cells were treated with control siRNA (Silencer Negative Control siRNA #2 (ThermoFisher Silencer Select, AM4613)) or KIF11 siRNA (antisense sequence: AUUGUCUUCAGGUCUUCAGtt; (ThermoFisher Silencer Select, 4390827)) for 24 hr before fixing or imaging.

### Generation & validation of RPE1 inducible cell lines

HILO-RMCE technology was used to engineer cell lines with one insertion of a KIF11-mEm sequence under control of a doxycycline-inducible promoter, as previously described (24–28). To establish the RMCE acceptor cell lines, ∼60% confluent RPE1 cell cultures were infected with serial dilutions of the pEM584 lentiviral stock. Cells were incubated for 48hr before the addition of selection media containing 20 μg/ml blasticidin. Next, individual clones were isolated by plating single cells via serial dilutions to a 96-well plate, and growing colonies. Clones were maintained in media with 10 μg/ml blasticidin.

To generate the RPE1-KIF11-mEm expressing lines, an RMCE donor plasmid was designed containing an siRNA-resistant KIF11-mEm construct (13) and a puromycin resistance gene flanked by loxP sites. The KIF11-mEm WT and KIF11-mEM Y211F constructs were made via QuikChange site directed mutagenesis (Agilent) to introduce the fluorescent tag, and Gibson Assembly to insert the KIF11-mEm tagged sequence into the RMCE acceptor vector (pEM7984). The KIF11-Y211E RMCE donor plasmid was generated via mutagenesis of the KIF11-mEm RMCE donor plasmid (GenScript).

RPE1 RMCE acceptor cell lines were plated in a 6-well plate at 150,000 cells/well with antibiotic-free media 24 hr prior to transfection. Cells were then co-transfected with a KIF11 RMCE donor plasmid and a pEM784 nl-Cre recombinase plasmid at a 1:10 wt/wt ratio. To transfect 2 wells of a 6-well plate, 1350ng of total DNA was mixed with 4.5uL of Lipofectamine LTX (ThermoFisher Scientific) reagent in 200uL of Opti-MEM (Life Technologies), with 100 uL of the mixture added to each well. Media was exchanged to remove LTX reagents a minimum of 2 hr post-transfection. 24 hr post-transfection, media containing 10 μg/ml puromycin (ThermoFisher Scientific, A11138-03) was added to each well to select for cells with successful recombination events. After 48 hr, the concentration was increased to 20 μg/ml puromycin for stronger selection. After 48 hr of strong selection, the concentration was decreased back to 5 μg/ml for maintaining the cell lines. Cells were pooled between the two transfected wells. Genomic DNA from each RPE1 inducible cell line was extracted (QIAmp DNA Blood Mini Kit Qiagen #51106) and sequenced through the Y211 PTM site (Eurofins) to verify correct incorporation of the desired KIF11 construct. Expression of KIF11-mEm constructs was induced with addition of 2 μg/ml doxycycline (Thermo Fisher Scientific #BP26531) to cell culture media for 24 hr. The same RPE1 accepter cell clonal line was used to generate all of the KIF11-mEm lines.

### Cell fixation and immunofluorescence

Cells were seeded on acid-treated 12-mm glass coverslips in a 24-well dish and fixed in ice-cold methanol (ThermoFisher Scientific) containing 1% paraformaldehyde (Electron Microscopy Sciences) for 10 minutes on ice and then washed 3 times for 5 minutes each in Tris-Buffered Saline (TBS; 150 mM NaCl, 50 mM Tris base, pH 7.4). Coverslips were blocked with 20% goat serum in antibody-diluting buffer (Abdil: TBS pH 7.4, 1% bovine serum albumin, 0.1% Triton-X, 0.1% sodium azide) for 1 hr at room temperature with shaking. After blocking, coverslips were washed two times for 5 minutes each in 1X TBS. Primary antibodies were diluted in Abdil: mouse anti-GFP (1:2,000, 0.5 μg/ml, ThermoFisher), rat anti-tubulin YL (1:1500, 0.5 μg/ml, Millipore), rabbit anti-KIF11 (1:1000, 0.2 μg/ml, NOVUS Biologicals). Coverslips were incubated with diluted primary antibodies for 1 hr at room temperature with shaking. Coverslips were then washed three times for 5 minutes each with 1X TBS, before incubation at room temperature with secondary antibodies against mouse, rat, and rabbit IgG conjugated to Alex Fluor 488, 594, 647 (1:500, 1 μg/ml, Molecular Probes). All secondary antibodies were diluted 1:500 in Abdil. After incubation with secondary antibodies, coverslips were washed three times in 1X TBS for 5 minutes each before mounting in Prolong Gold anti-fade mounting medium with DAPI (Invitrogen Molecular Probes, P36935).

### Mitotic cell phenotypes after expression of KIF11 constructs

RPE1 KIF11-mEm inducible cells were seeded on 12-mm coverslips in a 24-well dish. 24 hr later endogenous KIF11 was depleted using siRNA transfection, and 4-5 hr later, KIF11-mEm expression was induced by the addition of 2 μg/ml doxycycline for 24 hr (knockdown of endogenous KIF11, and replacement with KIF11-mEM). Cells were then fixed and stained for DNA, KIF11, GFP, and tubulin as described above. From random fields of view, the number of monopolar, bipolar, or disorganized mitotic spindles were counted in each condition. Only bipolar spindles in metaphase were included. Images were acquired from a minimum of 3 experimental replicates for each cell line. Total number of cells analyzed for each condition: Control KD: 83, Control KD (+KIF11-WT-mEM): 73, KIF11 KD: 161, KIF11 KD (+KIF11-WT-mEM): 84, Control KD (+KIF11-Y211E-mEm): 70, KIF11 KD (+KIF11-Y211E-mEm): 247, Control KD (+KIF11-Y211F-mEm): 38, KIF11 KD (+KIF11-Y211F-mEm): 63.

### KIF11-mEm expression level quantification in RPE1 inducible lines

KIF11-mEm and endogenous KIF11 expression level across the different inducible lines was quantified by immunofluorescence, using coverslips from the same sets of experiments used for the mitotic cell phenotypes. Cells were fixed and stained for KIF11 and GFP as described above. Images were acquired from a minimum of 3 experimental replicates, and analyzed using Image J. Images were background subtracted, and intensity was measured using a mask of the tubulin signal for each mitotic cell to only quantify GFP or KIF11 signal on mitotic spindles. Fluorescence intensities were normalized to the average intensity of the control + KIF11-mEm condition for each cell line.

### Live cell imaging: monastrol washout & pole to pole velocity analysis

RPE1-KIF11-mEm inducible lines were seeded in a glass-bottom 24-well plate with approximately 50,000 cells per well. 24 hr later, cells were treated with KIF11 siRNA and doxycycline as described for fixed cell experiments. Several hours after treatment with siRNA and doxycycline, monastrol was added to each well for overnight incubation at 100 μM. The next day, 4 hr prior to live cell imaging, the cell culture media was replaced with CO_2_ independent media (Gibco) containing 10% FBS, 2 µg/mL doxycycline, SiR Tubulin (Cytoskeleton, diluted 1:10,000) and 100µM monastrol. Mitotic cells arrested as monopoles were identified in a given well, and their locations recorded using the 60X objective and 647nm channel. Prior to imaging a given chamber, media with monastrol was aspirated and cells were washed 4 times with 500 µL of CO_2_ independent media containing 10% FBS and 2 μg/ml doxycyline. Immediately after washing out monastrol, imaging was initiated at one frame per 30 sec for the duration of pole separation (approximately 30 min). Spindle pole separation was determined by measuring the distance between spindle poles for each frame over the course of the movie using ImageJ/Fiji.

### Fluorescence after photobleaching (FRAP)

RPE1-KIF11-mEm inducible cell lines were seeded (approximately 55,000 cells) into 35mm glass-bottom plates (MatTek Corp). 24 hr later, cells were treated with KIF11 or control siRNAs in the morning, and treated with doxycycline 4-5 hr later, as described for the mitotic cell phenotype analysis. Cells were imaged 24 hr later. On the day of imaging, the media was changed to CO2-independent media supplemented with 10% FBS and 2 μg/ml doxycycline.

Cells were imaged for 5s at 250 ms intervals prior to photobleaching, photobleached for 200 ms using a 405nm laser, and imaged for 20 s at 250 ms intervals after photobleaching. A circular ROI with a 0.7 µm diameter was used to determine the photobleached spot, an unbleached standard elsewhere in the cell, and a background cell-free area. These 3 ROIs were imaged throughout the movie. For interphase cells, unbleached and bleached ROIs were placed at opposite ends of the cell within the cytoplasm. For mitotic cells with bipolar spindles, the unbleached and bleached ROIs were placed at opposite ends of the spindle. For monopolar spindles, the unbleached and bleached ROIs were placed as far apart from one another as possible on the monopole. Movies were background subtracted and corrected for photobleaching due to imaging acquisition using the NIS Analysis Software (Nikon). Half times (t_1/2_) were calculated by finding the time required for fluorescence to recover to 50% of its final intensity plateau.

### Live and fixed cell microscopy

Fixed cell images and live cell movies were acquired on a Ti-E inverted microscope (Nikon Instruments) or on a Ti-2E inverted microscope (Nikon Instruments) both driven by NIS Elements (Nikon Instruments). Images were captured using either a Clara cooled charge-coupled device (CCD) camera (Andor) or Prime BSI scientific complementary metal-oxide-semiconductor (sCMOS) camera (Teledyne Photometrics) with a Spectra-X light engine (Lumencore). Imaging was performed using the following Nikon objectives: Plan Apo 40 × 0.95 numerical aperture (NA), Plan Apo λ 60 × 1.42 NA, and APO 100 × 1.49 NA. For live cell-imaging, cells were imaged in CO_2_-independent media (Life Technologies #18045-088) supplemented with 10% FBS (Life Technologies #16000-044) within environmental chambers held at 37°C. FRAP experiments were performed using an inverted Eclipse Ti-2E microscope (Nikon) equipped with an iLas Ring-TIRF module and point-focused 405 nm laser driven by NIS Elements.

### MT gliding assays with TIRFM

For the mixed motor gliding experiments, a specialized flow cell was constructed utilizing a microscope slide in conjunction with a hydrophobic coverslip (Prepared via silane coating, 0.5% (v/v) Dichloro-Dimethyl-Silane in Trichloroethylene and rinsing with methanol via sonication, 15 mins, 3 times), securely held in place by double-sided adhesive tape as described previously (29). The cell was filled with penta-His antibodies sourced from Qiagen at a 200 μg/mL concentration. By combining 2.5 μL of these antibodies with 22.5 μL of PEM80, a final dilution of 20 μg/mL was achieved. Following a 10-minute absorption period, a casein solution (5.55 mg/mL casein in 35 mM Pipes, 5 mM MgSO4, 1 mM EGTA, and 0.5 mM EDTA) was applied for an additional 10 minutes to block the surface effectively.

Subsequently, motor proteins in the desired ratio of KIF11-WT: KIF11-mutants suspended in the motility buffer (80 mM Pipes (pH 6.9), 50 mM CH3CO2K, 4 mM MgSO4, 1 mM DTT, 1 mM EGTA, 10 μM paclitaxel, and 1 mg/mL casein buffer) were introduced into the chamber. The solution was incubated for 5 minutes to allow the motors to bind to the antibodies via their His tags specifically. The subsequent step involved the addition of 1 μL of fluorescent microtubules (MTs), comprised of 10% Cy3-tubulin (Puresoluble.com) and 90% non-fluorescent tubulin. The MTs were mixed in 50 μL of motility buffer containing 2mM ATP, along with an oxygen-scavenging system (250 μg/mL glucose oxidase, 30 μg/mL catalase, and 4.5 mg/mL glucose) before introducing them into the sample chamber.

To monitor the movements of the fluorescent MTs, they were excited with a 488 nm laser (Sapphire 488–500 CDRH, Coherent), imaged via a custom total internal reflection fluorescence (TIRF) microscope (Nikon 1.49NA, 100× TIRF objective), and recorded using a Photometrics QuantEM 512SC EMCCD camera at ∼5 frames per sec. For precise tracking of the fluorescent MTs, image processing was carried out in MATLAB to minimize shot noise and enhance image clarity. The intensity profile of each MT frame was analyzed using ImageJ, where a median filter (five points) was applied, followed by a Loess filter for smoothing. The tip positions of each microtubule (MT) were determined by analyzing the convergence of the intensity profiles from image #1 and image #200 in a time series TIFF stack, which included 200 time points for most cases (and 1,000 time points for pseudo-dimer KIF11-Y211E). The initial and final positions of the tips were utilized to calculate the gliding velocities. Depending on the specific experiment, gliding displacements for 15 to 32 individual MTs within the field of view were measured to estimate the average velocities.

## Acknowledgments

This work was supported by NIH R01GM130556 to JMM, SSR, SPG and JS; NIH R35 GM144133 to JS; a Department of Education Cellular, Molecular, and Biomedical Sciences Graduate Assistance in Areas of National Need (CMB-GAANN) fellowship to AA; and NIH F31AR074887 to AFT.

## Supplemental Figure Legends

**Figure S1. Double phospho-mimetic mutations in KIF11 alter force production and MT interaction dynamics**. (A) Schematic of KIF11 constructs used for in-vitro experiments. Amino acid numbers are provided in parentheses. (B) Example single molecule force traces from optical trapping experiments. (C-E) Force, velocity, and MT binding kinetics of single-molecule KIF11 pseudo-dimer constructs were measured using an optical trap. (C) Plot of peak forces measured for each of the indicated constructs. (D) Plot of MT bound times measured for each of the indicated constructs. (E) Plot of bead rebinding intervals measured for each of the indicated constructs. (C-E) Data points represent individual single-molecule measurements (n = 9 and 11 single-molecule beads for Y231E-Y125E (236 events) and Y211E-Y231E (117 events), respectively). Mean ± SEM is indicated for each data set. Data were compared using a student t-test (* p<0.05, *** p<0.001).

**Figure S2. Quantification of siRNA-mediated KIF11 KD in RPE1 cells, and of KIF11-mEm expression across RPE1 inducible cell lines. (A)** RPE1 cells were treated with KIF11 siRNA for 24 hr and then fixed and stained for KIF11 and tubulin. Using a mask of the tubulin signal, the KIF11 intensity on spindle microtubules was measured for each cell and normalized to the control KD. 24 hr treatment with KIF11 siRNA resulted in a 76% knockdown of endogenous KIF11. Control KD: 81 cells, KIF11 KD: 112 cells. **(B)** KIF11-mEm expression was quantified across the cell lines and conditions in Figure 3. GFP intensity was measured using a mask of the tubulin signal and normalized to control KD (no doxycycline).

**Figure S3. KIF11 turnover is not affected by phosphorylation on Y211 during interphase.** KIF11-mEm construct turnover was measured in interphase RPE1 cells using fluorescence recovery after photobleaching (FRAP). RPE1 cells were induced to express KIF11-mEm-WT, Y211E, or Y211F constructs with 24 hr doxycycline treatment, and either control or KIF11 siRNA. Using a 405 nm laser, cells were photobleached in a small region of interest, avoiding the nucleus and peripheral regions of the cell. Fluorescence intensity recovery over the following 20 sec was measured. **(A)** Fluorescence intensity traces after photobleaching. Traces have been corrected for photobleaching due to image acquisition. **(B)** Half times (t_1/2_) calculated for each KIF11-mEm construct. Cell counts for each cell line: **RPE1-KIF11-WT-mEM** - Control KD: 27 KIF11 KD: 21. **RPE1-KIF11-Y211E-mEm –** Control KD: 18, KIF11 KD: 20. **KIF11-Y211F-mEm –** Control KD: 20, KIF11 KD: 19. N = 3 experiments. Data were compared by running a one-way ANOVA with Šídák’s test for multiple comparisons: p-value style: <0.05 (*), <0.01 (**), <0.001 (***), and <0.0001 (****). If no significance is indicated, the result was not significant (>0.05).

**Movie S1: Spindle pole separation after monastrol washout.** Spindle polar separation was tracked in live movies of RPE1-KIF11-mEm expressing cells. Cells were induced to express KIF11-mEm constructs for 24 hr before live imaging with the addition of doxycycline and treated with control or KIF11 siRNA. Cells were also treated with monastrol overnight to inhibit KIF11 motors, collapse all spindles to monopoles, and arrest cells in mitosis. Spindles were labeled using SiR-tubulin several hours prior to imaging. Monastrol was washed out immediately before the start of imaging. **Top row:** Control KD with KIF11-WT (left), KIF11-Y211E (middle), and KIF11-Y211F (right). **Bottom row:** KIF11 KD with KIF11-WT (left), KIF11-Y211E (middle), and KIF11-Y211F (right).

**Movie S2: FRAP on mitotic cells expressing KIF11-mEm constructs.** KIF11-mEm construct turnover was measured in mitotic RPE1 cells using fluorescence recovery after photobleaching (FRAP). RPE1 cells were induced to express KIF11-mEm-WT, Y211E, or Y211F constructs with 24 hr doxycycline treatment, and either control or KIF11 siRNA. Using a 405 nm laser, cells were photobleached in a small region of interest on the spindle. **Top row**: KIF11-WT-mEm with Control KD (left) and KIF11 KD (right). **Middle row**: KIF11-Y211E-mEm with Control KD (left) or KIF11 KD (right). **Bottom row**: KIF11-Y211F-mEm with Control KD (left) or KIF11 KD (right).

## Notes

### Competing Interest Statement

The authors have declared no competing interest.

## References

1. A. Blangy, et al., Phosphorylation by p34cdc2 regulates spindle association of human Eg5, a kinesin-related motor essential for bipolar spindle formation in vivo. Cell 83, 1159–1169 (1995).

2. K. E. Sawin, T. J. Mitchison, Mutations in the kinesin-like protein Eg5 disrupting localization to the mitotic spindle. Proc. Natl. Acad. Sci. 92, 4289–4293 (1995).

3. D. J. Sharp, et al., The bipolar kinesin, KLP61F, cross-links microtubules within interpolar microtubule bundles of Drosophila embryonic mitotic spindles. J Cell Biology 144, 125–138 (1999).

4. J. Cahu, et al., Phosphorylation by Cdk1 Increases the Binding of Eg5 to Microtubules In Vitro and in Xenopus Egg Extract Spindles. PLoS ONE 3, e3936 (2008).

5. G. Goshima, R. D. Vale, Cell cycle-dependent dynamics and regulation of mitotic kinesins in Drosophila S2 cells. Mol Biol Cell 16, 3896–3907 (2005).

6. M. K. Chee, S. B. Haase, B-Cyclin/CDKs Regulate Mitotic Spindle Assembly by Phosphorylating Kinesins-5 in Budding Yeast. PLoS Genet. 6, e1000935 (2010).

7. J. D. Bishop, Z. Han, J. M. Schumacher, The Caenorhabditis elegans Aurora B Kinase AIR-2 Phosphorylates and Is Required for the Localization of a BimC Kinesin to Meiotic and Mitotic Spindles. Mol. Biol. Cell 16, 742–756 (2005).

8. O. Shapira, L. Gheber, Motile properties of the bi-directional kinesin-5 Cin8 are affected by phosphorylation in its motor domain. Sci. Rep. 6, 25597 (2016).

9. A. Goldstein, et al., Three Cdk1 sites in the kinesin-5 Cin8 catalytic domain coordinate motor localization and activity during anaphase. Cell. Mol. Life Sci. 74, 3395–3412 (2017).

10. R. Avunie-Masala, et al., Phospho-regulation of kinesin-5 during anaphase spindle elongation. J. Cell Sci. 124, 873–878 (2011).

11. J. M. Muretta, et al., A posttranslational modification of the mitotic kinesin Eg5 that enhances its mechanochemical coupling and alters its mitotic function. Proc. Natl. Acad. Sci. 115, E1779–E1788 (2018).

12. K. Garcia, J. Stumpff, T. Duncan, T. T. Su, Tyrosines in the kinesin-5 head domain are necessary for phosphorylation by Wee1 and for mitotic spindle integrity. Curr Biol 19, 1670–1676 (2009).

13. K. G. Bickel, et al., Src family kinase phosphorylation of the motor domain of the human kinesin-5, Eg5. Cytoskeleton 74, 317–330 (2017).

14. R. B. Irby, T. J. Yeatman, Role of Src expression and activation in human cancer. Oncogene 19, 5636–5642 (2000).

15. S. S. Rosenfeld, J. Xing, G. M. Jefferson, P. H. King, Docking and Rolling, a Model of How the Mitotic Motor Eg5 Works*. J. Biol. Chem. 280, 35684–35695 (2005).

16. J. M. Muretta, et al., The structural kinetics of switch-1 and the neck linker explain the functions of kinesin-1 and Eg5. Proc National Acad Sci 112, E6606–13 (2015).

17. S. Rice, et al., A structural change in the kinesin motor protein that drives motility. Nature 402, 778–784 (1999).

18. T. U. Mayer, et al., Small Molecule Inhibitor of Mitotic Spindle Bipolarity Identified in a Phenotype-Based Screen. Science 286, 971–974 (1999).

19. T. M. Kapoor, T. U. Mayer, M. L. Coughlin, T. J. Mitchison, Probing spindle assembly mechanisms with monastrol, a small molecule inhibitor of the mitotic kinesin, Eg5. The Journal of Cell Biology 150, 975–988 (2000).

20. B. J. Mann, P. Wadsworth, Kinesin-5 Regulation and Function in Mitosis. Trends Cell Biol 29, 66–79 (2018).

21. J. S. Waitzman, S. E. Rice, Mechanism and regulation of kinesin-5, an essential motor for the mitotic spindle. Biol Cell 106, 1–12 (2014).

22. K. J. Verhey, J. W. Hammond, Traffic control: regulation of kinesin motors. Nat Rev Mol Cell Bio 10, 765–777 (2009).

23. I. Nabti, et al., The ubiquitous microtubule-associated protein 4 (MAP4) controls organelle distribution by regulating the activity of the kinesin motor. Proc. Natl. Acad. Sci. United States Am. 119, e2206677119 (2022).

24. K. A. Queen, A. Cario, C. L. Berger, J. Stumpff, Modification of the neck-linker of KIF18A alters Microtubule subpopulation preference. Mol. Biol. Cell 35, ar3 (2024).

25. P. Khandelia, K. Yap, E. V. Makeyev, Streamlined platform for short hairpin RNA interference and transgenesis in cultured mammalian cells. Proc National Acad Sci 108, 12799–12804 (2011).

26. C. Marquis, et al., Chromosomally unstable tumor cells specifically require KIF18A for proliferation. Nat. Commun. 12, 1213 (2021).

27. A. F. Thompson, et al., Pathogenic mutations in the chromokinesin KIF22 disrupt anaphase chromosome segregation. eLife 11, e78653 (2022).

28. K. L. Schutt, et al., Identification of the KIF18A alpha-4 helix as a therapeutic target for chromosomally unstable tumor cells. Front. Mol. Biosci. 11, 1328077 (2024).

29. A. Goulet, et al., Comprehensive structural model of the mechanochemical cycle of a mitotic motor highlights molecular adaptations in the kinesin family. Proc National Acad Sci 111, 1837–1842 (2014).

