## Supplemental Figures for "A Phosphorylation Switch Governs KIF11’s Mechanical Output During Mitosis"

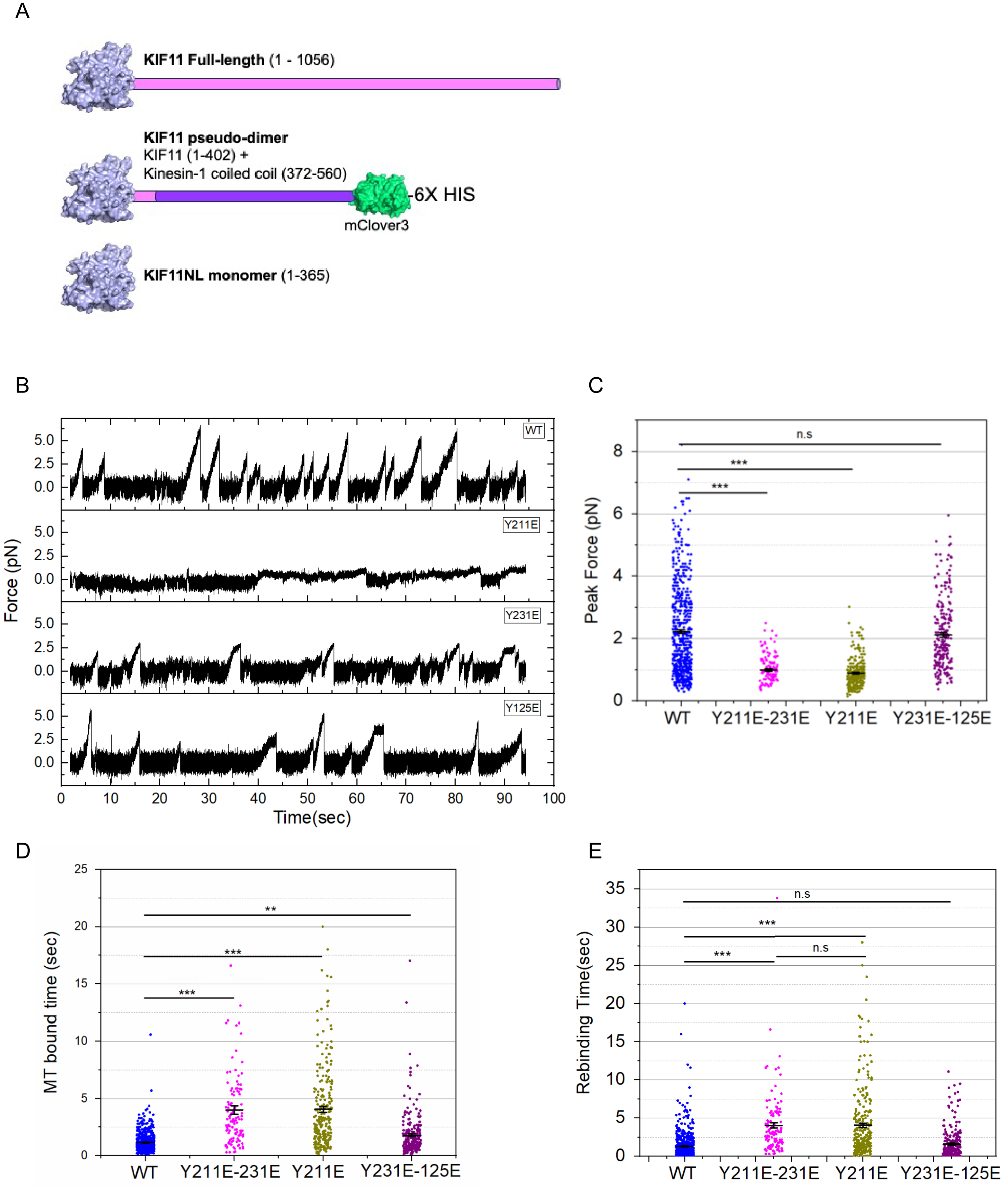

A
KIF11 Full-length (1 - 1056)
KIF11 pseudo-dimer
KIF11 (1-402) +
Kinesin-1 coiled coil (372-560)
-6X HIS
mClover3
KIF11NL monomer (1-365)
B
WT
Y211E
Y231E
Y125E
Force (pN)
Time(sec)
C
Peak Force (pN)
n.s
***
***
WT
Y211E-231E
Y211E
Y231E-125E
D
MT bound time (sec)
**
***
***
WT
Y211E-231E
Y211E
Y231E-125E
E
Rebinding Time(sec)
n.s
***
***
n.s
WT
Y211E-231E
Y211E
Y231E-125E

**Figure S1. Double phospho-mimetic mutations in KIF11 alter force production and MT interaction dynamics**. (A) Schematic of KIF11 constructs used for in-vitro experiments. Amino acid numbers are provided in parentheses. (B) Example single molecule force traces from optical trapping experiments. (C-E) Force, velocity, and MT binding kinetics of single-molecule KIF11 pseudo-dimer constructs were measured using an optical trap. (C) Plot of peak forces measured for each of the indicated constructs. (D) Plot of MT bound times measured for each of the indicated constructs. (E) Plot of bead rebinding intervals measured for each of the indicated constructs. (C-E) Data points represent individual single-molecule measurements (n = 9 and 11 single-molecule beads for Y231E-Y125E (236 events) and Y211E-Y231E (117 events), respectively). Mean ± SEM is indicated for each data set. Data were compared using a student t-test (* $p<0.05$, *** $p<0.001$).

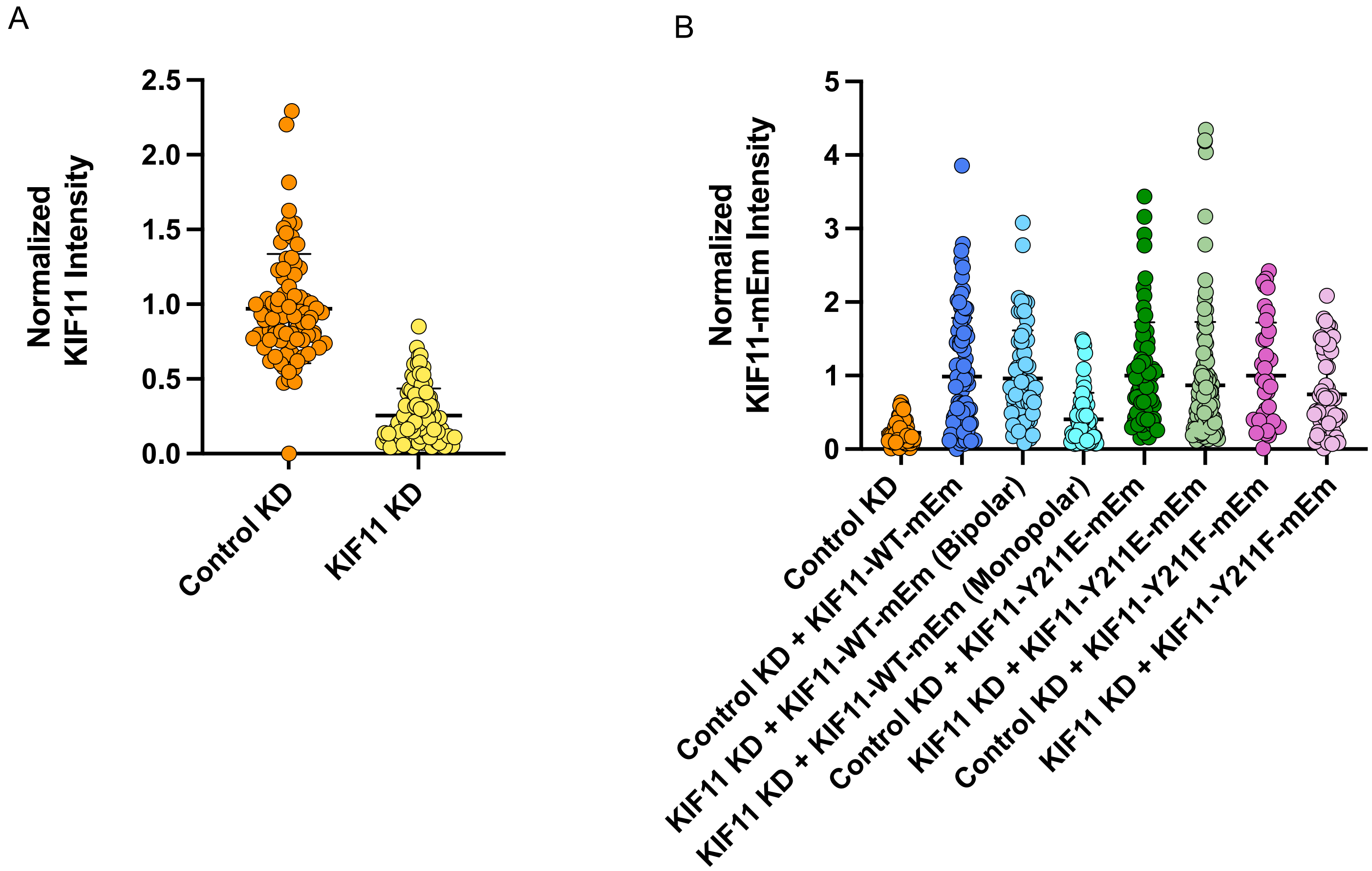


**Figure S2. Quantification of siRNA-mediated KIF11 KD in RPE1 cells, and of KIF11-mEm expression across RPE1 inducible cell lines**. (A) RPE1 cells were treated with KIF11 siRNA for 24 hr and then fixed and stained for KIF11 and tubulin. Using a mask of the tubulin signal, the KIF11 intensity on spindle microtubules was measured for each cell and normalized to the control KD. 24 hr treatment with KIF11 siRNA resulted in a 76% knockdown of endogenous KIF11. Control KD: 81 cells, KIF11 KD: 112 cells. (B) KIF11-mEm expression was quantified across the cell lines and conditions in Figure 3. GFP intensity was measured using a mask of the tubulin signal and normalized to control KD (no doxycycline).

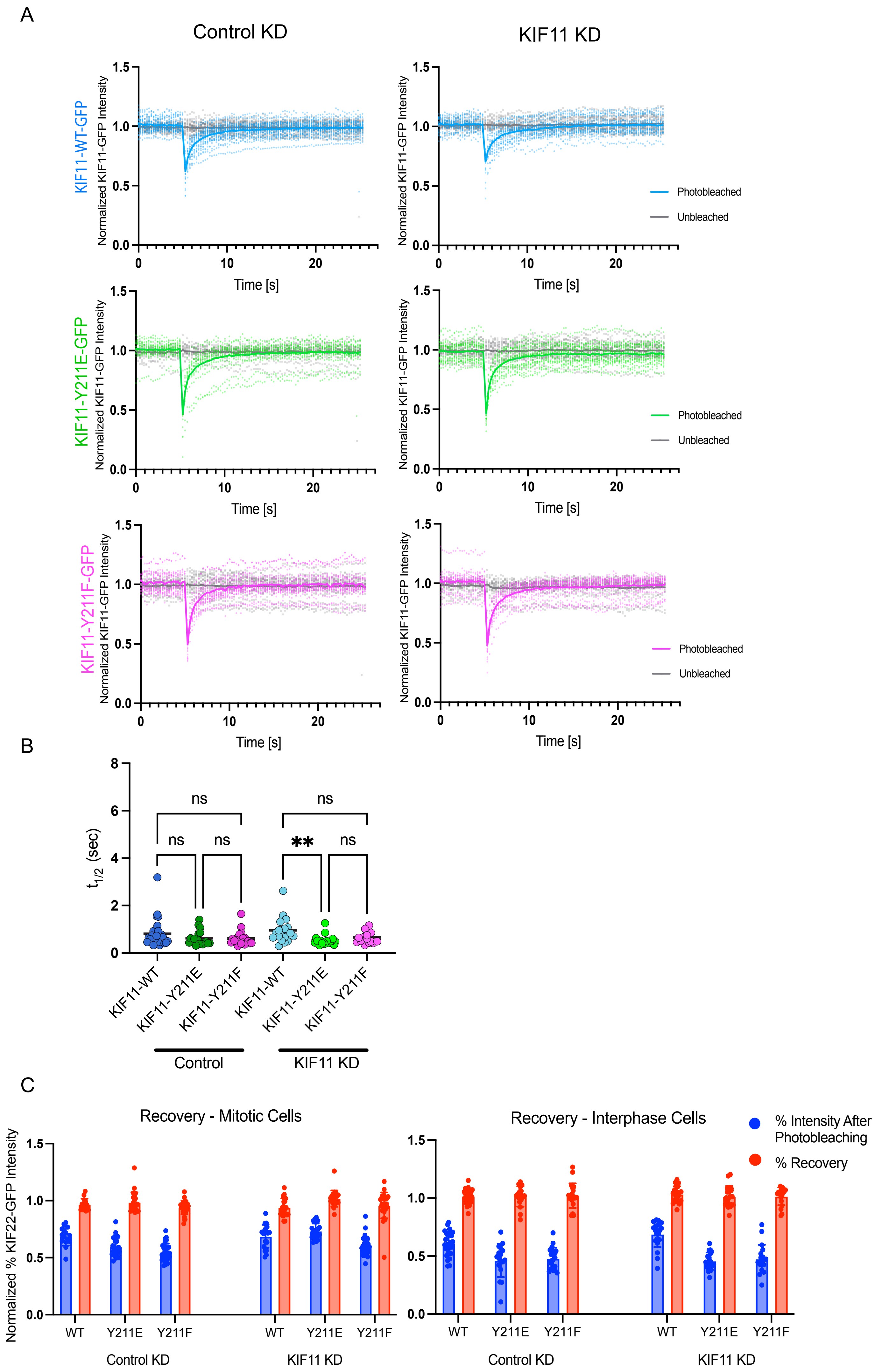
A
Control KD
KIF11 KD
KIF11-WT-GFP
KIF11-Y211E-GFP
KIF11-Y211F-GFP
Normalized KIF11-GFP Intensity
Time [s]
Photobleached
Unbleached
B
$t_{1/2}$ (sec)
ns
**
KIF11-WT
KIF11-Y211E
KIF11-Y211F
Control
KIF11 KD
C
Recovery - Mitotic Cells
Recovery - Interphase Cells
% Intensity After Photobleaching
% Recovery
Normalized % KIF22-GFP Intensity
WT
Y211E
Y211F
Control KD
KIF11 KD

**Figure S3. KIF11 turnover is not affected by phosphorylation on Y211 during interphase.** KIF11-mEm construct turnover was measured in interphase RPE1 cells using fluorescence recovery after photobleaching (FRAP). RPE1 cells were induced to express KIF11-mEm-WT, Y211E, or Y211F constructs with 24 hr doxycycline treatment, and either control or KIF11 siRNA. Using a 405 nm laser, cells were photobleached in a small region of interest, avoiding the nucleus and peripheral regions of the cell. Fluorescence intensity recovery over the following 20 sec was measured. **(A)** Fluorescence intensity traces after photobleaching. Traces have been corrected for photobleaching due to image acquisition. **(B)** Half times ($t_{1/2}$) calculated for each KIF11-mEm construct. Cell counts for each cell line: **RPE1-KIF11-WT-mEM** - Control KD: 27 KIF11 KD: 21. **RPE1-KIF11-Y211E-mEm –** Control KD: 18, KIF11 KD: 20. **KIF11-Y211F-mEm –** Control KD: 20, KIF11 KD: 19. N = 3 experiments. Data were compared by running a one-way ANOVA with Šídák's test for multiple comparisons: p-value style: <0.05 (*), <0.01 (**), <0.001 (***), and <0.0001 (****). If no significance is indicated, the result was not significant (>0.05).

**<u>Supplemental Movies</u>:**

**Movie S1: Spindle pole separation after monastrol washout.** Spindle polar separation was tracked in live movies of RPE1-KIF11-mEm expressing cells. Cells were induced to express KIF11-mEm constructs for 24 hr before live imaging with the addition of doxycycline and treated with control or KIF11 siRNA. Cells were also treated with monastrol overnight to inhibit KIF11 motors, collapse all spindles to monopoles, and arrest cells in mitosis. Spindles were labeled using SiR-tubulin several hours prior to imaging. Monastrol was washed out immediately before the start of imaging. **Top row:** Control KD with KIF11-WT (left), KIF11-Y211E (middle), and KIF11-Y211F (right). **Bottom row:** KIF11 KD with KIF11-WT (left), KIF11-Y211E (middle), and KIF11-Y211F (right).

**Movie S2: FRAP on mitotic cells expressing KIF11-mEm constructs.** KIF11-mEm construct turnover was measured in mitotic RPE1 cells using fluorescence recovery after photobleaching (FRAP). RPE1 cells were induced to express KIF11-mEm-WT, Y211E, or Y211F constructs with 24 hr doxycycline treatment, and either control or KIF11 siRNA. Using a 405 nm laser, cells were photobleached in a small region of interest on the spindle. **Top row**: KIF11-WT-mEm with Control KD (left) and KIF11 KD (right). **Middle row**: KIF11-Y211E-mEm with Control KD (left) or KIF11 KD (right). **Bottom row**: KIF11-Y211F-mEm with Control KD (left) or KIF11 KD (right).
